# High-throughput 3D imaging flow cytometry of adherent 3D cell cultures

**DOI:** 10.1101/2023.07.10.548361

**Authors:** Minato Yamashita, Miu Tamamitsu, Hiromi Kirisako, Yuki Goda, Xiaoyao Chen, Kazuki Hattori, Sadao Ota

## Abstract

Three-dimensional (3D) cell cultures are indispensable in recapitulating *in vivo* environments. Among many 3D culture methods, the strategy to culture adherent cells on hydrogel beads to form spheroid-like structures is powerful for maintaining high cell viability and functions through an efficient supply of nutrients and oxygen. However, high-throughput, scalable technologies for 3D imaging of individual cells cultured on the hydrogel scaffolds are lacking. This study reports the development of a high-throughput, scalable 3D imaging flow cytometry (3D-iFCM) platform for analyzing spheroid models on hydrogel beads. This platform is realized by integrating a single objective lens-based fluorescence light-sheet microscopy with a microfluidic device employing a combination of hydrodynamic and acoustofluidic focusing techniques. This integration enabled an unprecedentedly high-throughput, robust optofluidic 3D imaging, processing 513 cells s^-1^ and a total of more than 10^4^ cells within a minute. The large dataset obtained allows us to quantify and compare the nuclear morphology of adhering and suspended cells, revealing adhering cells have smaller nuclei with non-round surfaces. This platform’s high throughput, robustness, and precision for analyzing the morphology of subcellular compartments in 3D culture models holds promising potential for various biomedical analyses, including image-based phenotypic screening of drugs with spheroids or organoids.

## 1. Introduction

Three-dimensional (3D) cell culture models are invaluable *in vitro* platforms for biomedical research because they recapitulate *in vivo* complex environments better than two-dimensional (2D) culture methods^[1]^. For example, cancer organoids derived from patient samples mimic the original cancer pathophysiology more closely than the cell culture on a 2D surface^[2–4]^. This inherent capability of 3D culture to provide a more physiologically relevant context underscores the need for the development of suitable scaffolds that support cell proliferation, differentiation, and function. Beyond conventionally used hydrogel scaffolds such as Matrigel and collagen, intensive efforts have been made to engineer diverse 3D scaffold materials that emulate the physiological extracellular matrix. Resultantly, a variety of materials, ranging from hydrogel beads^[5,6]^, polymeric hard material^[7,8]^, and glass fiber^[9]^, have been utilized for 3D cell culture.

Among the array of 3D scaffold options, hydrogel beads serve as scalable and versatile platforms for conducting 3D cell culture^[10]^. The hydrogel bead-based culture offers a unique advantage in providing high cell yield at scale as the small beads, dispersed in the culture media, supply a larger surface area for cell adhesion compared to the 2D surface of well-bottoms^[11]^. In addition, the hydrogel bead-based system promotes cell fitness by ensuring an efficient supply of nutrients and oxygen^[12]^. Maintaining cell fitness at scale is crucial for biomedical applications such as the expansion of mesenchymal stem cells^[13,14]^ and human induced pluripotent stem cells^[15,16]^ for regenerative medicine and the mass-production of recombinant proteins^[17]^, antibodies^[18]^ and extracellular vesicles^[19]^.

To perform large-scale assays or screening for a cell population cultured in a 3D hydrogel bead-based system, high-throughput, scalable technologies for 3D imaging of individual cells are desirable^[20,21]^. However, the current golden standard tools for 3D imaging, such as confocal and light-sheet fluorescence microscopes, are often time-consuming^[22]^ and have scalability determined by the size of devices or wells to place cells in for imaging. A promising alternative optical analysis approach is flow cytometry, as it offers the capacity to process a large volume of cells through continuous flow-based scanning. However, conventional high-throughput flow cytometry and recently advanced 2D imaging flow cytometry (iFCM) techniques are typically employed for suspended cells and lack the capability of depth resolution. Thus, the existing flow cytometric methods fall short of fully characterizing individual adherent cells inhabiting the 3D space^[23]^. Several groups have developed 3D-iFCM techniques compatible with 3D culture models^[24–26]^, yet their throughput and scalability have not been quantitatively demonstrated, limiting their further applications.

In this work, we report the development of a high-throughput, robust, scalable 3D-iFCM platform for analyzing adherent 3D cell culture models. **Figure 1** depicts the schematic representation of our platform. The process begins with culturing adherent cells on microscale hydrogel beads to form uniformly sized spheroids on a large scale. We then captured 3D images of these spheroids by flowing them in a microfluidic channel, where a diagonal 2D cross-section of the channel is continuously imaged by a single objective lens-based fluorescence light-sheet imaging technique called oblique-plane microscopy (OPM)^[27,28]^. To ensure stable and high-speed flow-based parallel scanning of spheroids, we applied a combination of hydrodynamic and acoustofluidic focusing to the flowing spheroids within the microfluidic device. This approach confines the spheroid stream within a narrow semi-2D region, achieving uniform velocities of each spheroid passing through the optical section. Using this opto-acoustic-fluidic system, we realized a flow-based OPM imaging of the 3D cultured cells adherent to the hydrogel beads with unprecedented throughput and robustness, enabling scalable measurements. We record an analysis throughput of 37 spheroids and 513 cells per second. Furthermore, we performed a large-scale analysis of the 3D images of >10^4^ epithelial cells cultured on the beads (spheroids). We successfully retrieved the nuclear morphology of 13,358 cells comprising 954 spheroids, revealing that the nuclei of the cells adhering to the beads had smaller and more complex shapes, compared to when they were suspended in a solution.

**Figure 1.**
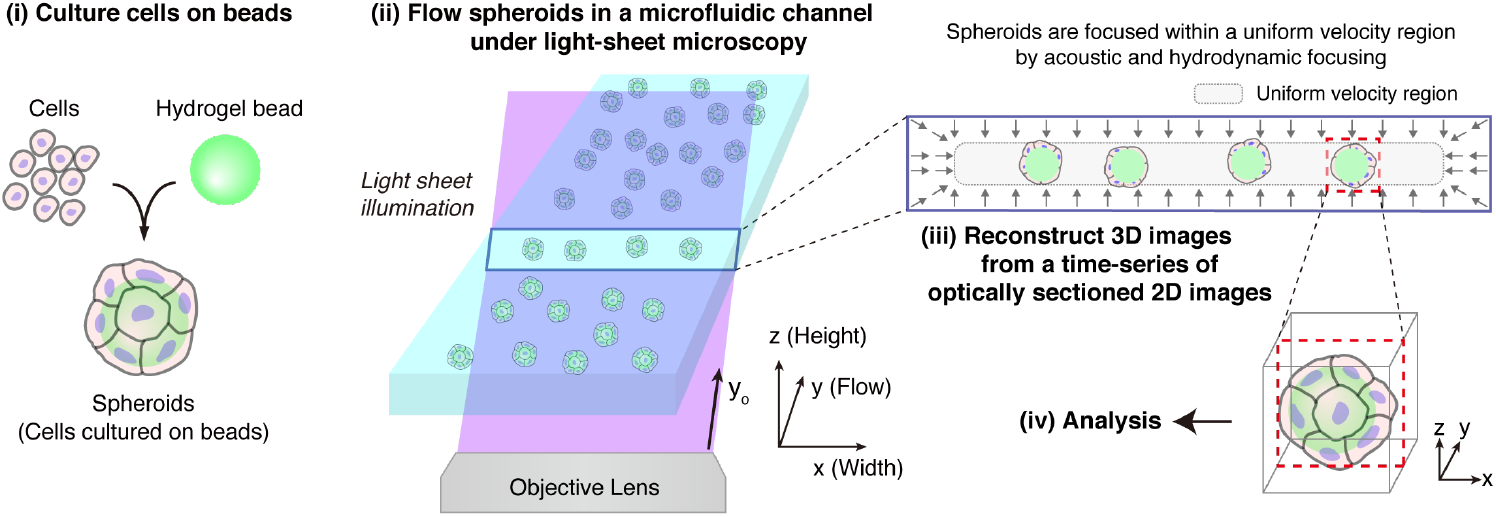
A scalable, high-throughput 3D imaging flow cytometry (3D-iFCM) platform for analyzing adherent 3D cell cultures. (i) First, detached adherent cells are incubated with alginate hydrogel beads so that multiple cells can attach to each bead. A spheroid-like structure forms when the cells proliferate and cover the surface of the beads. (ii) Then, we flow the spheroids in a microfluidic channel, where a diagonal 2D cross-section of the channel is continuously imaged by a single objective lens-based fluorescence light-sheet imaging technique. In the microchannel, we use a combination of acoustic and hydrodynamic focusing to confine the positions of the samples within a rectangular region, where the flow velocity is uniform (the gray area indicated in the upper right panel). (iii) Using this setup, we record a time series of optically sectioned 2D images of the focused spheroids and stack them to reconstruct 3D images, (iv) which are used to perform quantitative image analysis.

## 2. Results

### Focusing spheroids for high-throughput 3D imaging

In the experiment, we cultured Madin-Darby canine kidney (MDCK) cells on alginate hydrogel micro-beads that we prepared in advance using droplet microfluidics^[29]^. **Figure 2**a shows the phase-contrast and fluorescence images of the MDCK spheroids taken by a standard 2D microscope, in which green fluorescence is used to label the alginate hydrogel beads and cell membrane while blue fluorescence the cell nuclei. The images show that the MDCK cells adhere to and proliferate on the spherical hydrogel beads while the detailed 3D morphology of the cells cannot be resolved from these images.

**Figure 2.**
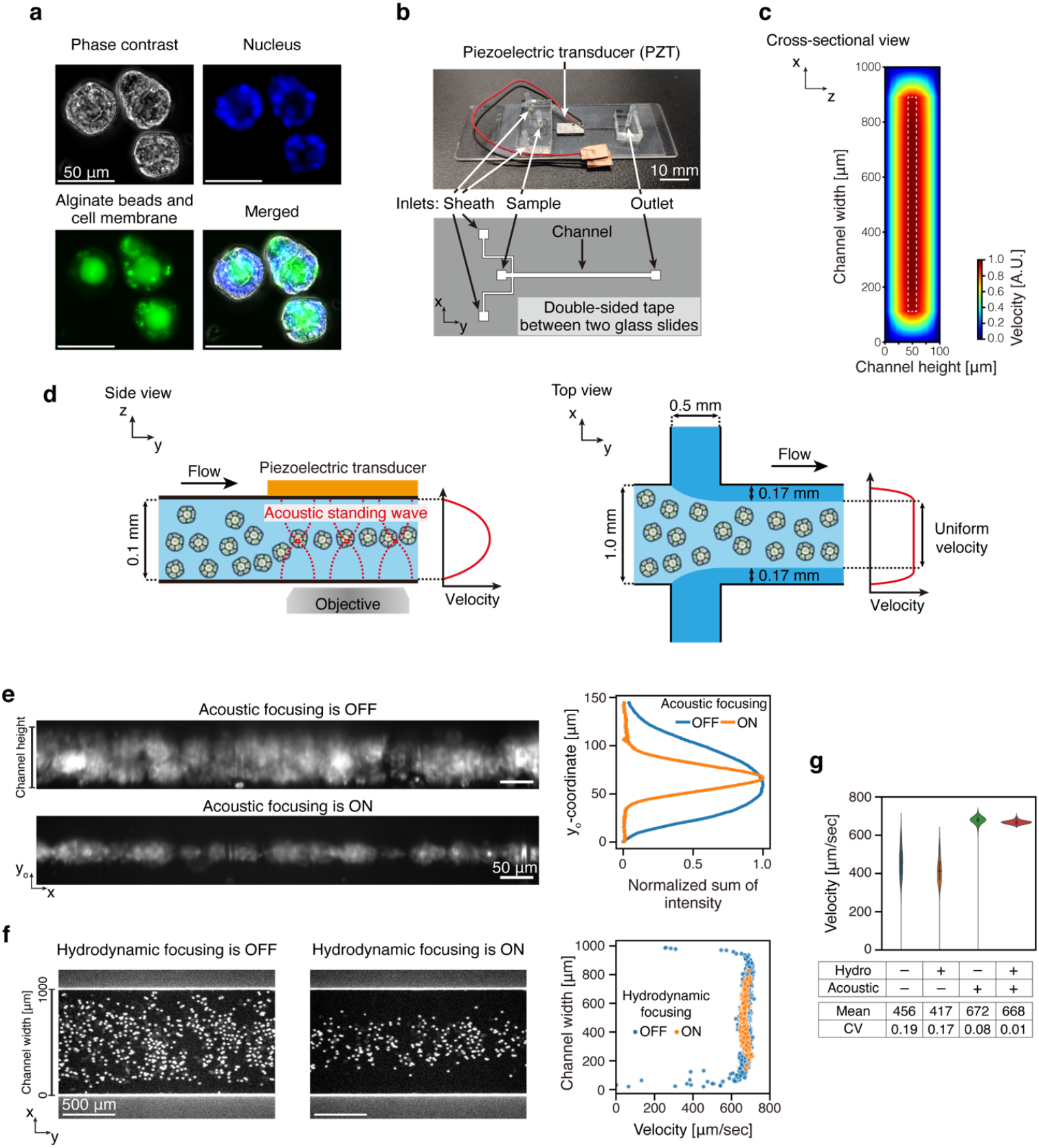
A combinatorial acoustic and hydrodynamic focusing approach to achieve uniform velocities of individual spheroids in a microchannel. (a) Representative phase-contrast and fluorescence images of the spheroids (MDCK cells cultured on alginate hydrogel beads). Blue: nucleus (Hoechst 33324); green: cell membrane (CellBrite Green) and bead (Alexa Fluor 488). Scale bar = 50 μm. (b) A photograph of our microfluidic device, consisting of a PZT and two glass slides bonded together by a microfabricated double-sided tape (top) that is designed to form a microchannel (bottom). (c) Numerical simulation of the velocity distribution inside the microfluidic channel. The rectangular region corresponds to the cross-section of the microfluidic channel. Inside the white-dotted rectangle, the flow velocity is mostly uniform. (d) Schematics of focusing techniques for the equalization of flow velocities of individual spheroids. Left: Acoustic focusing. Right: Hydrodynamic focusing. (e) Spheroid alignment in the y_o_-axis direction by acoustic focusing. Left: The intensity sums of 2D sectioned fluorescence images sequentially acquired for one second when acoustic focusing was off or on, respectively. Right: The normalized sum of the fluorescence intensity profile toward the channel height direction. The profile was calculated by summing the 2D sectioned fluorescence images, sequentially obtained for 30 seconds, and then integrating fluorescence intensities into the channel width direction. Scale bar = 50 μm. (f) Spheroid alignment by hydrodynamic focusing in the x-axis direction. Left and middle: Representative top-view 2D images of the flowing spheroids when hydrodynamic focusing was off or on, respectively. Right: Velocity distributions across the channel width. Scale bar = 500 μm. (g) Equalization of velocities of the spheroids by the two focusing methods. Velocity distributions in four conditions are shown in the violin plots, where acoustic focusing and hydrodynamic focusing are on or off, respectively.

Figure 2b shows a photograph of our microfluidic device and the design of the microfluidic channel. The device consists of two glass slides with custom-made double-sided tape in between them, creating a microfluidic channel with a width of 1 mm and a height of 100 μm. For acoustic focusing, the piezoelectric transducer (PZT) is attached to one of the outer glass surfaces of the device that does not face the objective lens. For hydrodynamic focusing, the device has one inlet for the sample and two inlets for the sheath fluid. To estimate the velocity distribution inside the microfluidic channel with a rectangular cross-section, we performed a numerical simulation assuming the Hagen–Poiseuille flow^[30]^ as shown in Figure 2c. The flow velocity increases toward the center of the channel along both the height and width directions and reaches 95% of the maximum value at ∼40 μm away from the top and bottom glasses and ∼100 μm away from the side walls. Therefore, the flow velocity of the spheroids is expected to be mostly uniform within the rectangular region (approximately 800 μm × 20 μm) indicated by the white dotted line (Figure 2e). To confine the flowing spheroids within the uniform velocity region, it is necessary to focus them along both the channel height and width directions. To this end, we adopt two focusing methods, acoustic focusing^[31,32]^ and hydrodynamic focusing^[33]^, as shown in the left and right panels in Figure 2d, respectively. In acoustic focusing, the PZT attached to the device induces acoustic standing waves that align the positions of spheroids to the middle of the channel in a height direction. Meanwhile, weak hydrodynamic focusing confines the position of the spheroids to the middle range of the channel in a width direction by two sheath flows applied from both sides. By optimizing flow rates, the widths of the two sheath flows and the sample flow are adjusted to be ∼0.17 mm and ∼0.66 mm, respectively. Collectively, the two focusing techniques allow us to achieve confinement of the spheroids within the uniform velocity region.

Figure 2e shows the successful alignment of spheroids by acoustic focusing. The images on the left side are the sums of 2D-sectioned green fluorescence images sequentially acquired for one second. When acoustic standing waves were off, the spheroids distributed in the height direction; however, when acoustic standing waves were on, the spheroids were aligned to the central region of the channel height. This acoustic focusing technique is robust enough for large-scale measurements (30 seconds; Movies S1, S2, Supporting Information). We further corroborated this focusing effect by calculating a line profile of the normalized sums of fluorescence intensity along the channel height (Figure 2e, right): we (1) summed the 2D sectioned green fluorescence images sequentially acquired for 30 seconds, then (2) summed the intensities of the summed images along the channel width direction, and (3) normalized the values by dividing them by the maximum value. The results quantitatively show the clear effect of acoustic focusing. Thus, our acoustic focusing method can focus flowing spheroids to the middle of channel height in a robust manner.

Next, we validated the effect of hydrodynamic focusing while acoustic focusing was on (Figure 2f). We sequentially obtained 2D top-view images of the spheroids on the hydrogel beads flowing in the channel with (left panel) or without (middle panel) hydrodynamic focusing for 9 seconds for each condition. When the hydrodynamic focusing was off, spheroids were distributed randomly across the channel width; when the hydrodynamic focusing was on, they gathered to the central region of the channel width. Then, we analyzed the motion of individual spheroids using a particle tracking method and quantified the average velocities of each spheroid (see Experimental Section for more details). The plot in the right panel shows the velocity distributions of the spheroids across the channel width. When hydrodynamic focusing was off, the velocities of the spheroids considerably varied because the velocities of the spheroids were low when they were flowing at regions < ∼100 μm from the walls, as expected from the simulation (Figure 2c). When hydrodynamic focusing was on, the spheroids were focused in the middle region of the channel width and exhibited almost uniform velocities. The width of the spheroid distribution, empirically confirmed, was ∼600 μm, which is included in the uniform velocity region predicted by the simulation (Figure 2c). Figure 2g summarizes the combinatorial focusing performance of acoustic and hydrodynamic methods in terms of the velocity distribution. The comparison shows that the combinatorial approach resulted in consistent and increased velocities of spheroids, where the mean was 667.9 μm s^-1^ and the coefficient of variation was 0.01 (Movie S3, S4, S5, and S6, Supporting Information). Overall, the combination of the acoustic and hydrodynamic focusing methods allowed us to focus spheroids within the central region where flow velocities are controlled uniformly (Movie S3 and S4, Supporting Information).

### High-throughput 3D imaging of spheroids

In **Figure 3**, we demonstrate the high-throughput 3D imaging capability of our platform. Figure 3a exemplifies the sequential and parallel acquisition of 2D optical sections of multiple spheroids by the camera. Blue fluorescence images show the cell nuclei and green ones show the alginate beads and cell membrane. We measured the section images at the frame rate of 200 Hz and display them with the frame interval of 10 milliseconds, which is equivalent to every two frames. These time series of images visualize the sequence of 2D cross-sections of nuclei, beads, and cell membranes, captured while spheroids flow through the light-sheet illumination. Figure 3b shows a reconstructed 3D image obtained within one second, wherein the blue and green fluorescence images are merged. To calculate the throughput of our system with these 3D images, we counted the number of spheroids and the number of cells by the number of nuclei contained inside the spheroids. The detailed procedure of the image analysis is explained in the next subsection (**Figure 4**a) and Experimental Section. As a result, the measured throughput was calculated as 37 spheroids s^-1^ and 513 cells s^-1^.

**Figure 3.**
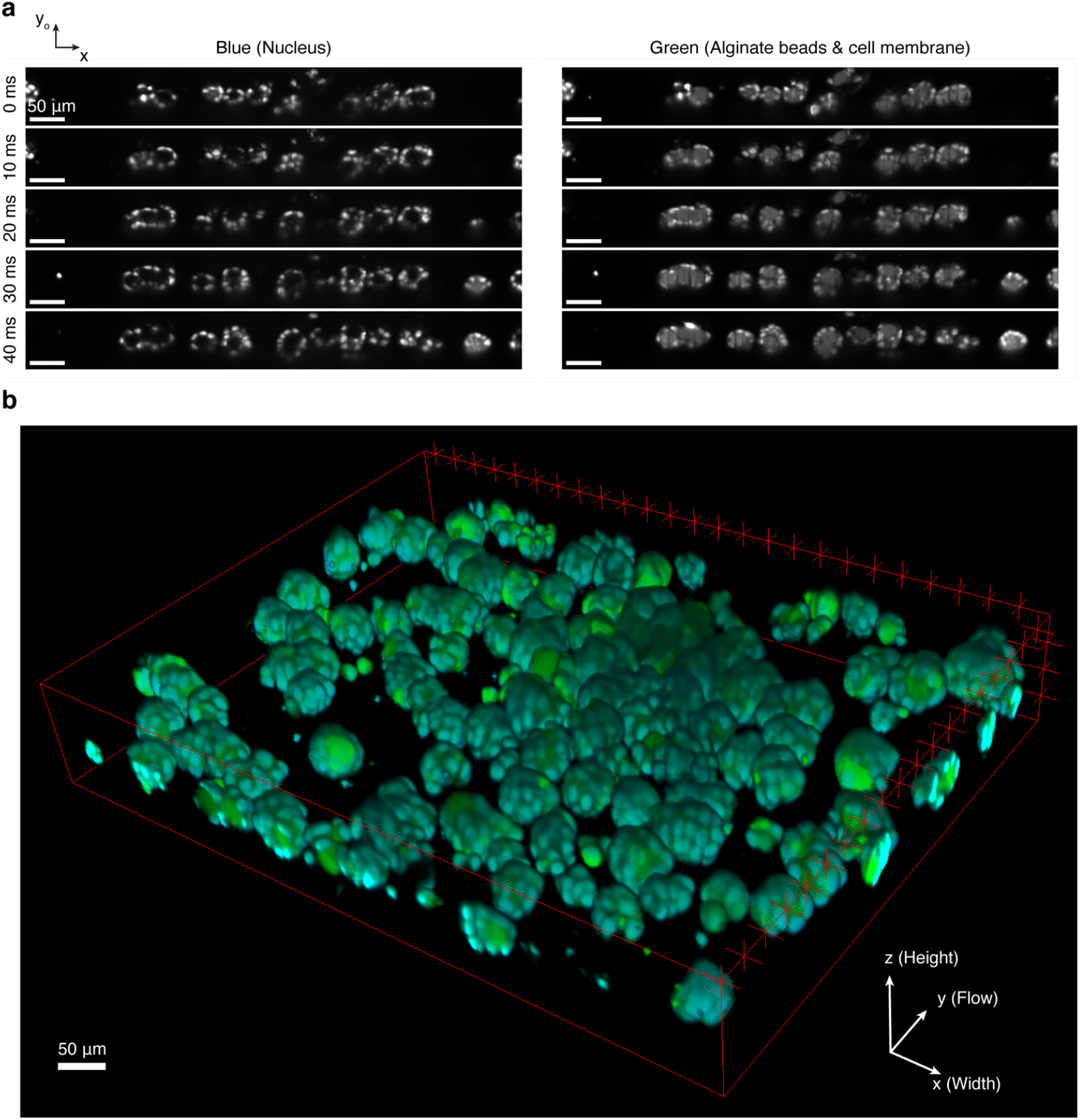
High-throughput 3D imaging of the spheroids on hydrogel beads. (a) Sequential and parallel acquisition of 2D optical sections of multiple spheroids. The camera was operated at 200 frames per second. The images are displayed with a frame interval of 10 milliseconds (every two frames). Blue: nucleus (Hoechst 33342); green: cell membrane (CellBrite Green) and bead (Alexa Fluor 488). Scale bar = 50 μm. (b) A 3D image reconstructed from the stack of the 2D images sequentially acquired in one second. The numbers of spheroids and cells in the spheroids are 37 and 513, respectively.

**Figure 4.**
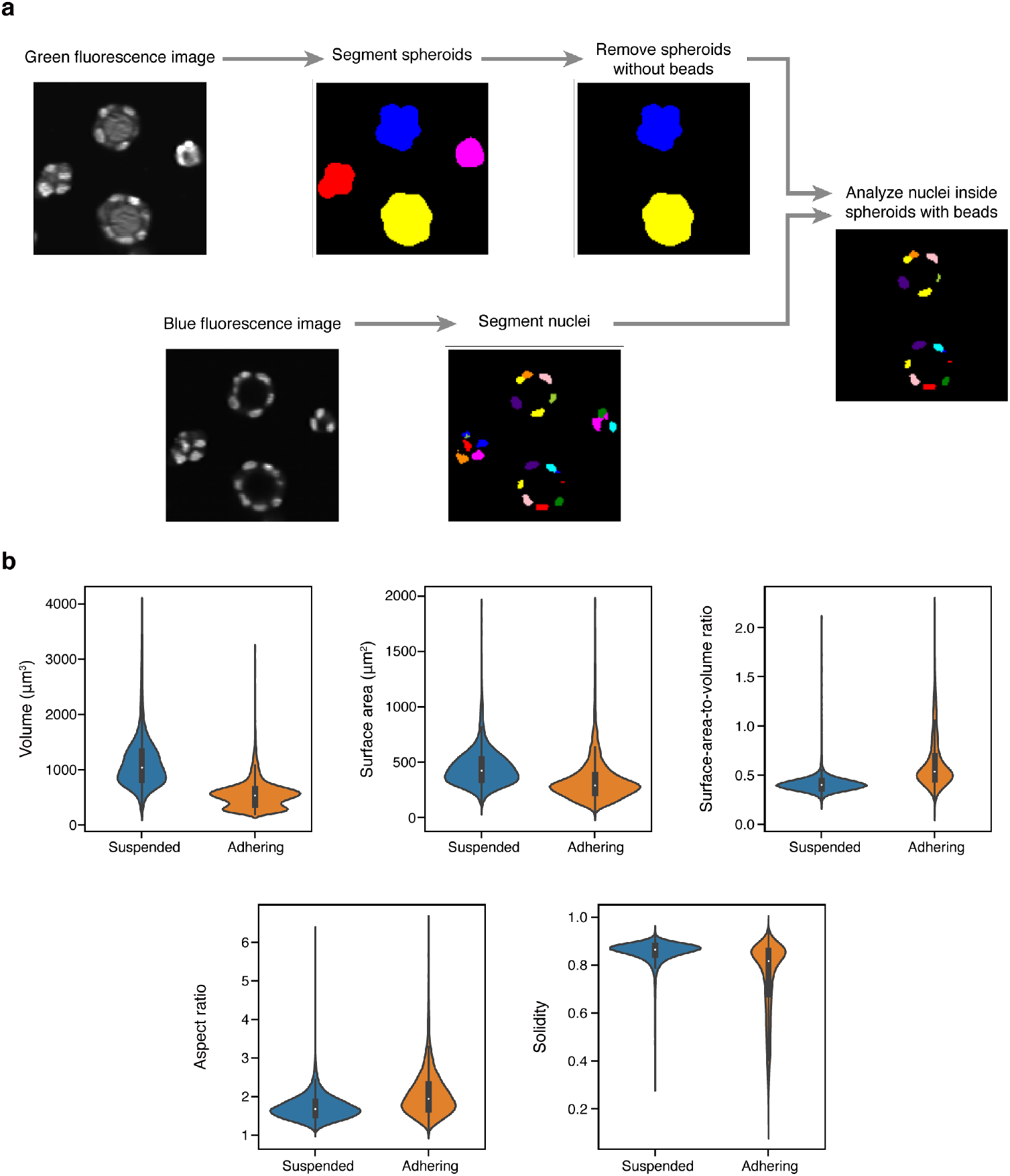
Comparative morphological analysis of suspended and adhering MDCK cells. (a) The procedure of image analysis for deriving nuclear morphological characteristics from the spheroids. First, spheroids and nuclei are segmented separately using green and blue fluorescence images, respectively. Specifically, spheroid masks were created by segmenting the regions surrounded by the cell membranes while excluding spheroids without beads inside. Then, from only nuclei located inside the spheroid masks, we extracted the morphological properties. (b) A side-by-side comparison of the nucleic morphology between MDCK cells adhering to beads and those suspended in a solution. We analyzed key morphological attributes, including volume, surface area, surface-area-to-volume ratio, aspect ratio, and solidity. The distributions of these properties are displayed using violin plots (n=13,358 and 27,160 for adhering and suspended states, respectively).

### Revealing the 3D nucleic morphology of adhering cells on beads on a large scale

As a biological demonstration of our robust, scalable platform, we performed the 3D-iFCM-based comparative analysis of nuclear morphologies of adhering and suspended states of MDCK cells (Figure 4). To this end, we prepared two samples: MDCK spheroids (adhering group), wherein the cells adhere to alginate beads, and suspended MDCK cells (suspended group) as a control. Using the 3D-iFCM platform, we individually measured the spheroids and the suspended cells at the frame rate of 300 Hz for 90 and 30 seconds, respectively. We then analyzed the reconstructed 3D images and quantified the nuclear morphological features of 13,358 cells (954 spheroids) and 27,160 cells for the adhering group and the suspended group, respectively. Figure 4a shows the image analysis procedure for extracting the morphologies of nuclei in spheroids.

Importantly, we selected only the cells cultured on the hydrogel beads, and thus mere cell aggregates (i.e., cells not cultured on the beads) were excluded from the subsequent analyses (see Experimental Section for details). For suspended MDCK cells, we analyzed all nuclei that were successfully segmented. As key morphological characteristics, we retrieved five morphological features of the nucleus: volume, surface area, surface-area-to-volume ratio, aspect ratio, and solidity^[34]^. Figure 4b summarizes the analysis result revealing differences in the nuclear morphology between the adhering and suspended groups (see also Figure S1, Supporting Information). Specifically, by analyzing more than 10^4^ cells in each group, we found that cells in the adhering group have smaller volumes and surface area of the nuclei, reflecting the smaller nuclei. Also, the nuclei of the adhering group showed a higher aspect ratio and a higher surface-area-to-volume ratio, indicating that the nuclear shape was non-spherical and more complex than the nuclei of the suspended group. This is likely because cells in spheroids were densely cultured on the beads and compacted by the adjacent cells. This observation is consistent with the fact that the adhering group exhibited lower solidity values, suggesting a higher presence of concavities in the nuclei. These results demonstrate that our 3D-iFCM can profile the organelle structure of adhering cells at high throughput on a large scale, which cannot be achieved by flow cytometry using detached adherent cells.

## 3. Discussion

To our knowledge, our platform has enabled the fastest and most scalable 3D-iFCM for 3D cultures. Yet, there is still room for improvement in the performance of our platform including throughput and scalability. First, the throughput of the platform is in trade-off with the spatial resolution. For example, by adjusting the magnification rate of the imaging system such that the sampling spatial resolution on the image sensor (which is currently ∼0.3 μm) is matched to the optical diffraction limit (which is ∼0.6 and ∼1 μm in x and y_o_ dimensions, respectively), we can double the imaging speed and hence the throughput since the frame rate of our camera is inversely proportional to the number of rows of the frame. Meanwhile, our current platform has another spatial resolution of ∼3.3 μm in the y dimension (i.e., along the flow) when it is operated at the frame rate of 200 Hz. Although this is slightly larger than the optical diffraction limit of ∼1 μm in this dimension, we are still able to quantify various morphological parameters of single cells. Therefore, if the spatial resolutions in the other (i.e., x and y_o_) dimensions can be sacrificed to be matched to our current y spatial resolution, we can further reduce the magnification rate to increase the camera frame rate and hence throughput by another factor of 5.5. With this reduced optical magnification, the field of view can be increased to the limit determined by the aberration-corrected range of the objective lens, which is ∼1.3 mm for our current objective lens, leading to another increase in the throughput by a factor of 1.73. Collectively, an order of magnitude improvement in the throughput can be expected, potentially leading to a throughput of ∼700 spheroids s^-1^ (currently 37 spheroids s^-1^). Second, the scalability can be expanded up to the limit determined by the data storage capacity, provided that the sample flow is stable. For example, with our current platform, the data spooling speed is 0.5 GB s^-1^ with each image-acquisition module (i.e., each fluorescence channel) operating at 200 Hz with a frame format of 16 bits and 512 × 2560 pixels. Thus, with a 2-TB high-speed data storage for each image-acquisition module, ∼1.5 × 10^5^ spheroids (assuming the current throughput of 37 spheroids s^-1^) can be measured within ∼1.1 hours with the same frame format. Moreover, with the reduced optical magnification, a ∼12 times reduction in the data size and hence an increase in scalability can be expected, potentially leading to a scalability of ∼1.8 × 10^7^ spheroids in an hour.

The capability of our platform to accurately analyze intracellular structures at high resolution and speed can be valuable for numerous biomedical studies focusing on adherent cells. Unlike conventional flow cytometry, our cell culture system leverages hydrogel beads to maintain adherent phenotypes while ensuring high throughput and scalability. While we analyzed adherent-specific nuclear morphologies in Figure 4, other subcellular attributes, such as the localization of proteins^[35]^ and co-localizations of proteins and organelles^[36,37]^ are also feasible measurement targets. For example, we can examine the polarization of MDCK cells on beads by quantifying the localized expression of the apical and basal markers^[38]^. Since the loss of apico-basal polarity can trigger various diseases including polycystic kidney disease, our system can be used to screen for perturbations that restore the polarity, which may lead to the development of novel therapeutics^[39,40]^. Compared to confocal or light-sheet microscopy without flowing samples, typically used for assessing subcellular properties including apico-basal polarity, our method offers superior throughput and scalability and will be, therefore, an effective platform for drug discovery^[41–43]^.

Furthermore, the potential applications of our platform can be extended to other 3D-culture settings, such as cancer spheroids and organoids. Cancer spheroids, self-assembled spherical cell aggregates, recapitulate the complex microenvironments in the cancer tissue^[20]^ and are, therefore, useful for drug screening^[44]^. Organoids, tissue-like miniaturized structures composed of multiple differentiated cell types, are useful for disease modeling and developmental biology studies^[45,46]^. Due to the heterogeneous nature of cell populations inside these 3D culture models, 3D imaging is crucial to fully characterize individual cells. Meanwhile, throughput and scalability pose significant challenges for these applications as comparing many conditions is desired for drug screening or analyses of patient-derived samples^[20]^. Our high-throughput and scalable platform may address these issues effectively. Nonetheless, we should further evaluate our platform using larger models since commonly used spheroids and organoids are usually 200 to 1,000 μm in diameter^[47]^ Implementing optical clearing methods^[48]^, non-diffracting and self-healing excitation beams^[49,50]^, and/or multi-view imaging approach^[51]^ for light-sheet microscopy may help us perform imaging of such large and highly-scattering biological samples.

## 4. Conclusions

In this study, we have engineered a high-throughput, robust, scalable 3D-iFCM platform for analyzing 3D cell culture models by integrating optofluidic parallel 3D light-sheet fluorescence microscopy with acoustic and hydrodynamic focusing techniques. We demonstrated the platform’s performance by 3D imaging of spheroids comprising MDCK cells on hydrogel-based at a throughput of 37 spheroids per second (513 cells s^-1^). We also showed the utility of this platform for quantitatively studying subcellular structures by extracting and comparing morphological features of the nucleus (n=13,358) in adhering cells and those (n=27,160) in suspended cells. The versatility of this method opens new avenues for various biological studies and biomedical applications, including image-based drug and phenotypic screening for spheroids and organoids.

## 5. Experimental Section

### 3D parallel light-sheet microscope

The design of the optical setup is shown in Figure S2 in the Supporting Information. For fluorescence excitation, two continuous-wave lasers are used with wavelengths of 405 nm (Stradus 405-250, Vortran) or 488 nm (OBIS LX 488 nm 150 mW Laser, Coherent). The laser beams are combined by a dichroic mirror, shaped into light sheets using a Powel lens (LOCP-8.9R05-1.0, Laserline Optics), and delivered to the sample through a series of 4f relaying optics. At the sample plane, the light sheets are ∼750 μm wide with an average power of 39 and 22 mW, respectively. The microscope is based on axial plane optical microscopy^[52,53]^ in which a 20×, NA-0.75, dry-type lens (UplanSApo20x, Olympus) is used for the primary and remote objective lenses. Two wide-field tube lenses (SWTLU-C, Olympus) are implemented between the primary and remote objective lenses to perform remote focusing. After the remote objective lens, two optical filters (T495lpxr-UF3 and T565lpxr-UF3, Chroma) are inserted to separate the fluorescence into two distinct spectral channels corresponding to the two excitation wavelengths. In the downstream path of each spectral channel, a tube lens (TTL200, Thorlabs) and a camera (Zyla 5.5, Andor) are implemented to capture the 2D optical section of the microfluidic channel. The total magnification rate of the microscope is ∼22.2, giving a sampling spatial resolution of ∼0.3 μm determined by the 6.5-μm pixel size of the camera. Each camera was operated at a frame rate of 200 Hz with 512 × 2560 pixels (Figure 2e, Figure 3a, b) or 300 Hz with 300 × 2560 pixels (Figure 4b) in 16 bits. The data are transferred to the computer via the CameraLink interface.

### Microfluidic conditions

The microfluidic device and the design of the microfluidic channel are shown in Figure 2b. A double-sided tape with the microfluidic channel of the design sticks two glasses. The device has three inlets: the central inlet is for the sample and the others are for the sheath fluid. Each of the three flows was introduced to the microfluidic channel with an independent syringe pump (Harvard Apparatus) via a PEEK tube. The flow rates were 2 μL min^-1^ for the sample and 0.5 μL min^-1^ for each sheath fluid, resulting in a total flow rate of 3 μL min^-1^. When the sheath flow was not used, the flow rate for the sample was 3 μL min^-1^. The inside of the device and the tube for the sample were coated with 0.5% Pluronic F-127 (Biotium, #59005) for more than one hour before injecting spheroids.

A piezoelectric transducer (PZT) with a resonance frequency of 3.75 MHz (3.75Z5 × 10R-SYX (C-213), Fuji Ceramics) was attached to the upper glass of the device. Applying a sine-wave signal with a peak-to-peak amplitude of 3.0 V amplified 10 times by an amplifier (HSA4101, NF Corporation) and a frequency of 7.5 MHz generated acoustic standing waves.

### Preparation of the alginate hydrogel beads

To enable fluorescence detection of the alginate hydrogel beads, we first labeled alginate by cross-linking Alexa Fluor 488 Hydrazide (Thermo Fisher Scientific, #A10436) with sodium alginate (IL-6, KIMICA). Then, 2% labeled alginate solution was mixed with 2% peptide-coupled alginate solution (NOVATACH LVM GRGDSP, Novamatrix, #4270701) at a ratio of 1:3. The alginate solution was further mixed with a solution of 50 mM calcium chloride and 50 mM EDTA at a ratio of 1:1.

Droplets were generated with a flow-focusing device using Droplet Generator oil for EvaGreen (Bio-Rad, #1864112) as a continuous flow. After collecting the intended number of droplets, gelation of the sodium alginate was induced by adding 0.05% acetic acid to the oil to release the calcium ion inside the droplets by decreasing the pH. Emulsions were then broken by adding HFE-7200 (3M) containing 20% 1H, 1H, 2H, and 2H perfluoro-1-octanol (Wako, #324-90642), and the hydrogel beads with a typical diameter of 40 μm were collected in a custom-made buffer composed of 10 mM Tris (pH8.0), 137 mM NaCl, 2.7 mM KCl and 1.8 mM CaCl_2_. This buffer is called the alginate buffer, hereafter.

### Cell culture on the alginate hydrogel beads

An epithelial cell line, Madin-Darby canine kidney (MDCK), was used for all experiments. MDCK cells were provided by the RIKEN BRC through the National Bio-Resource Project of MEXT, Japan. MDCK cells were maintained in DMEM-high glucose (Wako, #043-30085) containing 10% FBS (Sigma, #F7524) and 2 mM L-alanyl-L-glutamine (Wako, #016-21841) in a 5% CO2 atmosphere at 37°C. First, 1.35 × 10^6^ suspended MDCK cells were mixed with 1.35 × 10^5^ alginate beads in 1 mL of culture medium and stably incubated in a 6-well plate at 37 °C for two hours. Then, the cell suspension was diluted 10 times by adding 9 mL culture medium and replaced in a cell-repellent 10 cm dish (Greiner, #657970). The dish was continuously shaken reciprocally at 70 rpm for five days on a shaker (Taitec, CS-LR) in a CO2 incubator.

### Cell preparation for measurements

Collected spheroids were fixed with a paraformaldehyde (PFA) solution containing calcium chloride for 20 minutes. This PFA solution was prepared by mixing 4% PFA in PBS with 1M CaCl_2_ at a ratio of 9:1 and incubating it for more than 20 minutes. It was centrifuged at 20,000 × g for 5 min before use to remove the precipitate. After washing the spheroids with the alginate buffer twice, the cell membrane was stained with CellBrite Green Cytoplasmic Membrane Dye (1:200, Biotium, #30021) for 1 hour, and the nucleus was stained with Hoechst 33342 (20 μg/mL, Thermo Fisher Scientific, #H1399) for 15 min. Suspended cells were prepared by trypsinizing the cells cultured in a 10 cm dish, fixed, and stained as mentioned above. The fixed spheroids and suspended cells were filtered through a 50-μm cell strainer (pluriStrainer 50 μm, pluriSelect Life Science, #43-50050-03) and a 40-μm cell strainer (mini Cell Strainers II, Hitech, #HT-AMS-04002) to remove large aggregates, respectively. Then they were diluted before experiments so that the concentrations of the spheroids containing only one bead inside and the suspended cells were 5 × 10^5^ spheroids mL^-1^ and 5 × 10^6^ cells mL^-1^, respectively. All procedures were performed at room temperature.

### Numerical simulation of flow velocity distribution

The numerical simulation was performed with a custom-made code written in Python3. The velocity field for the Hagen–Poiseuille flow in a rectangular channel is written as

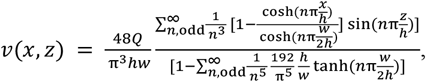

where *x* and *z* are coordinates inside the microfluidic channel with the dimension of *w* × *h* and *Q* is flow rate^[30]^. The values of *w* and *h* were fixed to the real values of the used microfluidic channel, namely *w*=1 and *h*=0.1. Fourier sums of this equation were approximately calculated using a library sympy (version 1.11.1). By repeatedly calculating velocities across a cross-section of a microfluidic channel, a velocity distribution inside the channel was obtained.

### Image analysis environment

All image analyses were conducted with custom-made codes written in Python3. Basic image processing and quantification were performed using the Python library Scikit-image (version 21.0). Also, we adopted two libraries, namely pyclesperanto_prototype (version 0.24.1) and cellpose^[54]^ (version 2.1.1), for the segmentation of the spheroid and nucleus, respectively. All reconstruction processes and analyses of 3D images were conducted on a workstation equipped with a GPU, GeForce GTX TITAN X (NVIDIA).

### Quantification of velocities of spheroids

The velocity distributions of the spheroids inside the microfluidic channel were measured using a standard 2D microscope. A temporal series of epi-fluorescence images of the flowing spheroids were acquired with an inverted microscope (ECLIPSE Ti-U, Nikon) equipped with a 4× objective lens (CFI Plan Fluor DL 4XF, Nikon), a GFP filter set (Chroma), and a CMOS camera (ASI1600MM Pro, ZWO) operating at 40 frames per second. Each frame is an image covering the width of the microfluidic channel. The flow and focusing conditions were the same as the 3D measurements. However, cells were not stained to simplify subsequent image analyses, and only fluorescence from beads was used to define flowing spheroids. Measurements were conducted for 9 seconds for each condition (i.e., acoustic focusing and hydrodynamic focusing are on or off, respectively). The images were analyzed with a Python library, trackpy (version 0.5.0). Beads were detected in each frame, and their movements were tracked throughout the time-series measurement by connecting the positions of the same beads between two adjacent frames. The beads whose tracks were found in less than 3 frames were removed from subsequent analyses. Finally, the velocities of the beads were obtained by averaging all the velocities of the same beads between two adjacent frames.

### Morphological analysis

The nuclei of the cells are segmented based on the blue (nuclei staining) fluorescence images, while spheroids are segmented based on the green (membrane and bead staining) fluorescence images. Volume [μm^3^] and solidity of the spheroids were calculated from the segmented image of the spheroids. To eliminate spheroids containing no beads inside, spheroids with < 30,000 μm^3^ in volume and < 0.8 in solidity were removed, and the others were retained for the subsequent analyses. Figure S3a shows histograms of volume and solidity calculated from a part of the data. It was visually confirmed that these thresholds worked almost successfully (Figure S3b). Next, we quantified the nuclear properties of cells inside the retained spheroids. Volume [μm^3^], surface area [μm^2^], surface-area-to-volume ratio, aspect ratio, and solidity were chosen as the morphological features of nuclei. For analyzing suspended cells, after segmenting nuclei, all detected nuclei were used for the analysis.

## Acknowledgments

We thank Dr. Masashi Ugawa for his support and discussions and Yuka Mori for her technical support for this project. This work was supported by JST CREST grant number JPMJCR19H1 and JSPS KAKENHI grant numbers JP21H04636, JP21H00416, and JP22K12797, Japan. This work was also supported by Tateisi Science and Technology Foundation, the Noguchi Institute, the Naito Foundation, the Uehara Memorial Foundation, the White Rock Foundation, the Canon Foundation, Senri Life Science Foundation, the SECOM Science and Technology Foundation, Takeda Science Foundation, the Cell Science Research Foundation, the Japan Society for the Study of Obesity, and the UTEC-UTokyo FSI Research Grant Program.

## Data Availability Statement

Data and codes are available on BioStudies (https://www.ebi.ac.uk/biostudies/studies/XXX) and GitHub (https://github.com/solabtokyo/Yamashita_et_al_2023), respectively.

## Supporting Information

**Figure S1.**
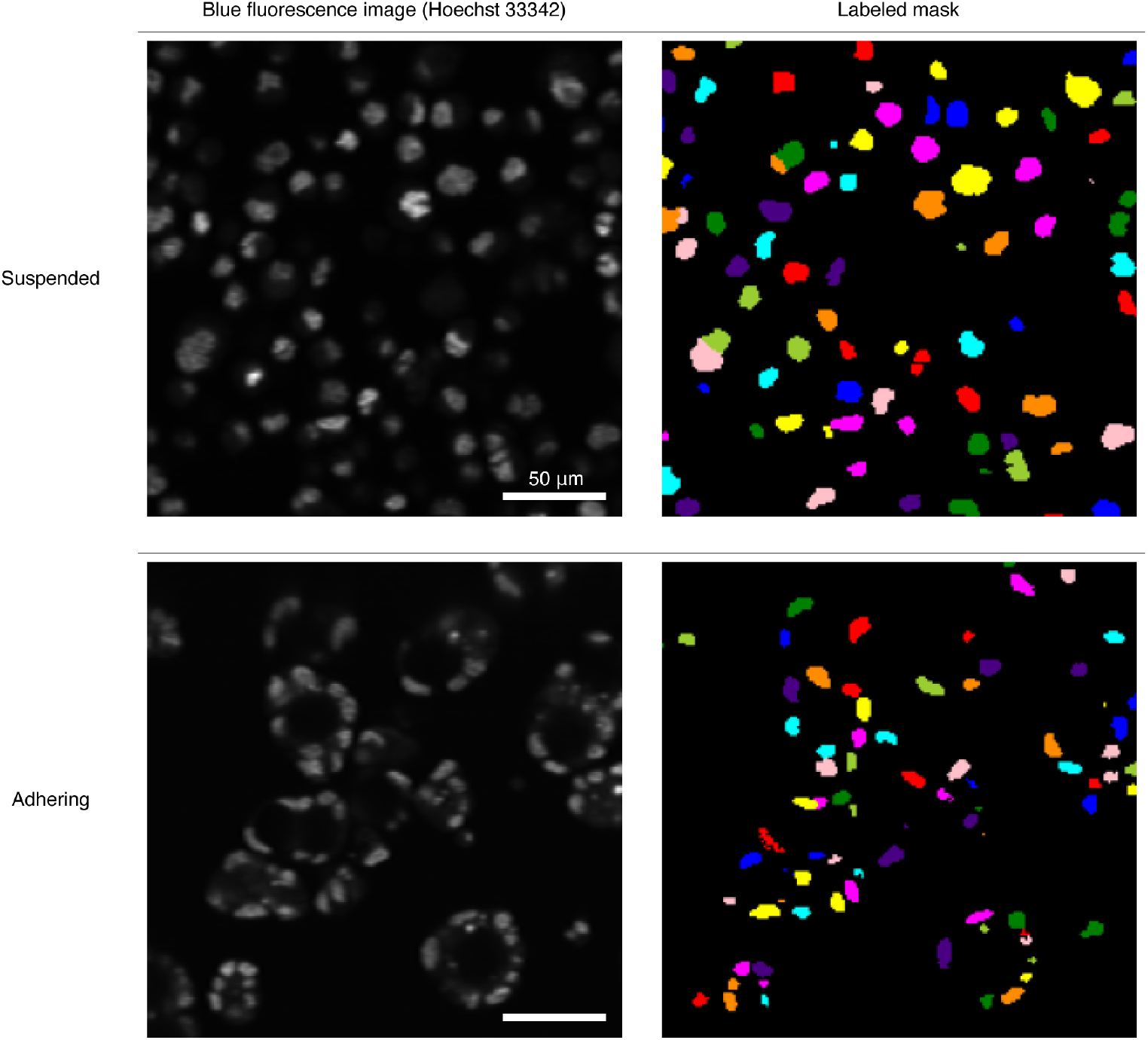
Examples of nuclear segmentation of suspended cells and adhering cells on beads. The left images show the nuclei stained with Hoechst 33342; the right images show the results of segmentation. Each segmented region is colored according to the labels assigned to the region. All images are z-slices from the 3D images.

**Figure S2.**
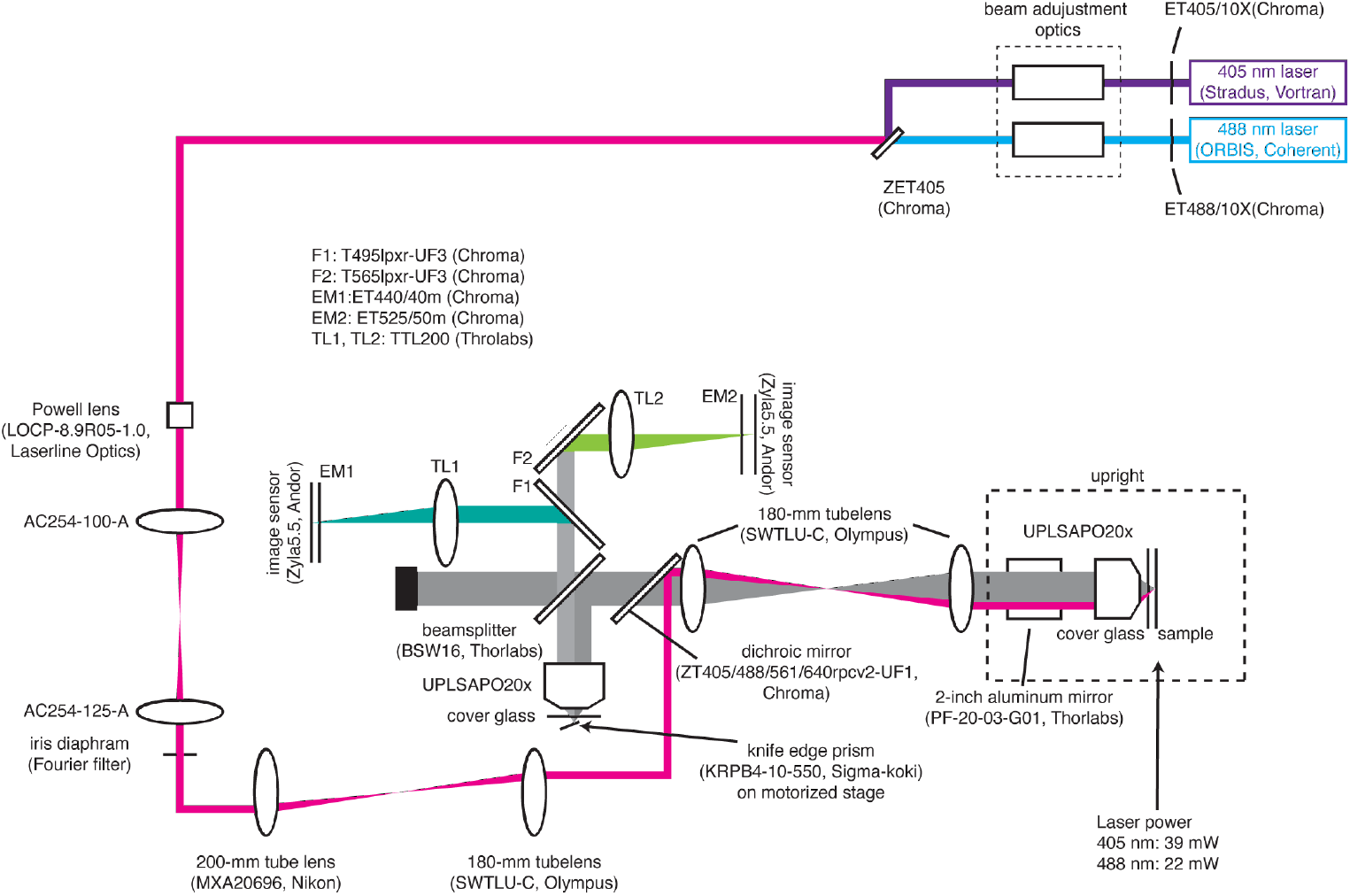
Detailed optical setup of the 3D parallel light-sheet microscope.

**Figure S3.**
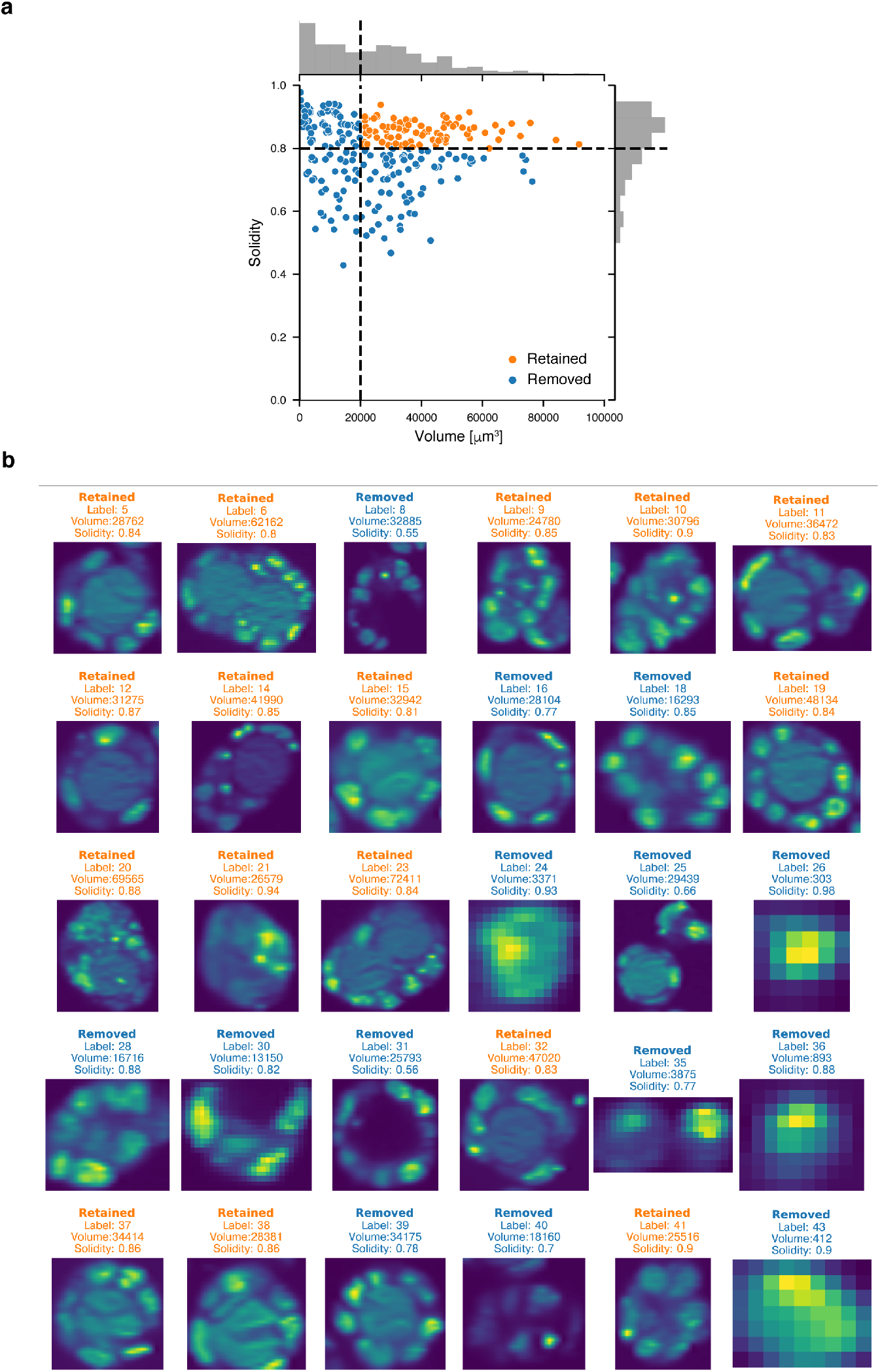
Representative results in the selection process of spheroids containing beads. (a) A scatter plot of the volume and solidity of the spheroids – both spheroids with or without beads are included. The black dashed lines indicate the threshold values (volume: 20,000; solidity: 0.8) for the selection of spheroids for the subsequent analyses; dots corresponding to the selected spheroids are shown in orange. n=294. (b) 30 representative z-slice images from the 294 segmented spheroids. On the top of each image, the result of selection (retained or removed) for the subsequent analyses, the label of the spheroid, and the values of volume and solidity were shown (Orange: Retained, Blue: Removed).

Supplemental Movie 1 and 2 are cross-sectional green fluorescence images of spheroids at the xy_o_ plane. Cell membrane: CellBrite Green, bead: Alexa Fluor 488.

**Supplemental Movie 1**. Images when acoustic focusing and hydrodynamic focusing are off.

**Supplemental Movie 2**. Images when acoustic focusing is on and hydrodynamic focusing is off.

Supplemental Movie 3-6 are top-view green fluorescence images of spheroids at the xy plane. Bead: Alexa Fluor 488.

**Supplemental Movie 3**. Images when acoustic focusing and hydrodynamic focusing are off.

**Supplemental Movie 4**. Images when acoustic focusing is on and hydrodynamic focusing is off.

**Supplemental Movie 5**. Images when acoustic focusing is off and hydrodynamic focusing is on.

**Supplemental Movie 6**. Images when acoustic focusing and hydrodynamic focusing are on

